# Integrated structural model of the palladin-actin complex using XL-MS, docking, NMR, and SAXS

**DOI:** 10.1101/2024.08.25.609580

**Authors:** Rachel Sargent, David H. Liu, Rahul Yadav, Drew Glennenmeier, Colby Bradford, Noely Urbina, Moriah R. Beck

**Author notes:** Corresponding Author: Moriah R. Beck, 1845 Fairmount Street, CB 51, Wichita, Kansas 67260.

## Abstract

Palladin is an actin binding protein that accelerates actin polymerization and is linked to metastasis of several types of cancer. Previously, three lysine residues in an immunoglobulin-like domain of palladin have been identified as essential for actin binding. However, it is still unknown where palladin binds to F-actin. Evidence that palladin binds to the sides of actin filaments to facilitate branching is supported by our previous study showing that palladin was able to compensate for Arp2/3 in the formation of *Listeria* actin comet tails. Here, we used chemical crosslinking to covalently link palladin and F-actin residues based on spatial proximity. Samples were then enzymatically digested, separated by liquid chromatography, and analyzed by tandem mass spectrometry. Peptides containing the crosslinks and specific residues involved were then identified for input to HADDOCK docking server to model the most likely binding conformation. Small angle X-ray scattering was used to provide further insight into palladin flexibility and the binding interface, and NMR spectra identified potential interactions between palladin’s Ig domains. Our final structural model of the F-actin:palladin complex revealed how palladin interacts with and stabilizes F-actin at the interface between two actin monomers. Three actin residues that were identified in this study also appear commonly in the actin binding interface with other proteins such as myotilin, myosin, and tropomodulin. An accurate structural representation of the complex between palladin and actin extends our understanding of palladin’s role in promoting cancer metastasis through regulation of actin dynamics.

**Significance:** In this study we have combined various advanced structural biology techniques to provide the first comprehensive model of the palladin-actin complex. Considering palladin’s role in cancer cell metastasis, this structure could be useful in screening and developing chemotherapeutic agents that target this interaction and prevent cancer cell metastasis.

## Introduction

Actin is a cytoskeletal protein that is abundant in eukaryotic cells and is crucial for cell shape and motility. Actin performs its functions through the dynamic reorganization of monomers into filaments and vice versa through polymerization or depolymerization, respectively. Many proteins are required to modulate actin’s functions in the cytoskeleton and muscle, and these proteins are known as actin binding proteins (ABPs). Palladin is a widely expressed ABP that is known to bind to actin and promote complex structures known as crosslinks or bundles. Palladin is critical during times of high cell motility such as embryonic development (26) and wound healing (38; 4) and is also known to be upregulated in several types of metastatic cancers including breast (13), pancreatic (14), adult gliomas (28), renal cell carcinoma (17), and non-small cell lung cancer (40).

There are over 150 known ABPs that bind, bundle, cap, sever, and further regulate actin within the cell. Some actin binding motifs in ABPs have been identified: calponin homology (CH) (12), Wiskott-Aldrich syndrome homology (WH2) (31), gelsolin homology (49), actin depolymerizing factor (ADF-H) domains (33), and RPEL domains (containing RPxxxEL repeats) (16). However, several immunoglobulin-like (Ig) domain containing proteins, like palladin and its protein family members, are also known to bind to actin despite not containing these recognized actin-binding motifs. Although the longest isoform of palladin contains five Ig domains, only the Ig3 domain of palladin directly binds to actin—both the monomeric actin (G-actin) (1) and F-actin forms (9). Interestingly, although the Ig4 domain does not bind to actin, the tandem Ig3-Ig4 (Ig3-4) construct binds and bundles F-actin more efficiently than Ig3 alone (9) perhaps due in part to a unique, elongated linker region between the Ig domains. Therefore, our experiments presented here focus on the Ig3 and the tandem Ig3-4 domains to explore possible structural reasons for this enhancement in binding.

Palladin has two solvent exposed basic patches on the surface of the Ig3 structure (PDB 2LQR *Mus musculus*, PDB 2DM2 *Homo sapiens*), and a previous study has identified several lysine residues that are critical for binding to negatively charged actin filaments (3). While the isolated Ig3 and Ig4 (PDB 2DM3 *H. sapiens*) domain structures have been solved by solution state NMR, no structural information is available to discern how the tandem Ig3-4 construct is oriented or where the Ig3 domain binds to actin filaments. Interestingly, a previous study of palladin’s role in actin comet tail formation of *Listeria monocytogenes* showed that the ubiquitously expressed isoform of palladin (containing the Ig3, Ig4, and Ig5 domains) was able to compensate for Arp2/3’s role in actin branching (8) suggesting that palladin is likely to bind to the sides of actin filaments.

To elucidate the molecular mechanism of palladin interaction with F-actin and the role of this interface in actin dynamics, we used an integrative structural approach involving crosslinking mass spectrometry (XL-MS), molecular docking, and small angle X-ray scattering (SAXS) to build the first structural model of the palladin:F-actin complex. Chemical crosslinking coupled with mass spectrometry has become an increasingly popular way of identifying bound protein interfaces and residues involved in binding (32). Covalent chemical crosslinkers vary in composition but share a common format: an active group on either end that binds to specific amino acids connected by a spacer arm. We utilized several chemical crosslinkers in this study with varying head groups and spacer arm lengths. Crosslinked samples of 1:1 actin to palladin complexes were trypsinized, subjected to liquid chromatography (LC), and assessed by tandem mass spectrometry (MS/MS). Interprotein linkages were identified and used as restraints to dock protein structures and generate a bound palladin:F-actin model using HADDOCK software. Docked structures were further refined and the highest scoring models were identified by a combination of variables including HADDOCK score and Xwalk score as discussed below. SAXS data is also presented that indicates a high degree of flexibility between Ig3 and Ig4 in the tandem construct and provides the first information regarding the orientation of these two domains in solution. Based on our results, we present the first structural model of palladin’s side-binding interaction with actin and identify actin residues in the interface that are commonly involved in the binding of other ABPs to actin as well. Considering palladin’s role in cancer cell metastasis, structural insights into the interactions between these two proteins could be useful in screening and developing chemotherapeutic agents that target this interaction and prevent cancer cell metastasis

## Results

### Integrative structural model of palladin:F-actin complex

To validate and identify potential new contacts between F-actin and palladin, we performed cross-linking coupled to mass spectrometry (XL-MS) experiments on Ig3 and tandem Ig3-4 domains of palladin bound to F-actin using a variety of crosslinking agents (BS3, DSSO, DFBNB, and DMTMM). Polymerized actin was incubated with various concentrations of the crosslinker and Ig3 or Ig3-4 before the reaction was quenched and the samples were run on an SDS-PAGE gel (Figure 1A). Bands at the molecular weight for a palladin:actin heterodimer were excised, trypsinized, and analyzed by LC-MS/MS. The zero-length crosslinker 4-(4,6-dimethoxy-1,3,5-triazin-2-yl)-4-methylmorpholiniumchloride (DMTMM) which links carboxylic acids and amine groups yielded the most consistent cross-links as expected given the likely electrostatic nature of the interaction between palladin and F-actin (Figure 1B). Other crosslinking agents (BS3, DSSO, and DFBNB) also produced crosslinked species but yielded few and inconsistent crosslinks (data not shown), so the DMTMM results were used as the restraints to guide macromolecular docking. As expected, in both the Ig3 and Ig3-4 results, most crosslinks were identified between lysine residues (K13, K36, K46, and K51) in the Ig3 domain and aspartic or glutamic acid residues on actin (D24, D25, D51, E99, and E100) (Figure 1C-E, Table 1). Many of these actin residues have also been found in the binding interface between actin and other ABPs (Table 2). Consistent with previous studies finding no binding between F-actin and the isolated Ig4 domain, and no crosslinks were detected from Ig4 residues. Interestingly, although no interaction between actin and the isolated linker region has been discovered previously (9), results from the Ig3-4 XL-MS indicate that several residues at the end of the linker region are in close proximity with F-actin. In contrast to the crosslinks from Ig3 to F-actin, these link actin residue K328 to four negatively charged residues within the Ig3-4 linker, **D**_127_SG**D**_130_**E**_131_N**E**_133_ (Figure 1F). While the crosslinks to the linker region could not be utilized in our current modeling experiments due to lack of any structural information for this linker or tandem domain, these results could indicate an undiscovered function for this linker region that warrants further investigation.

**Figure 1:**
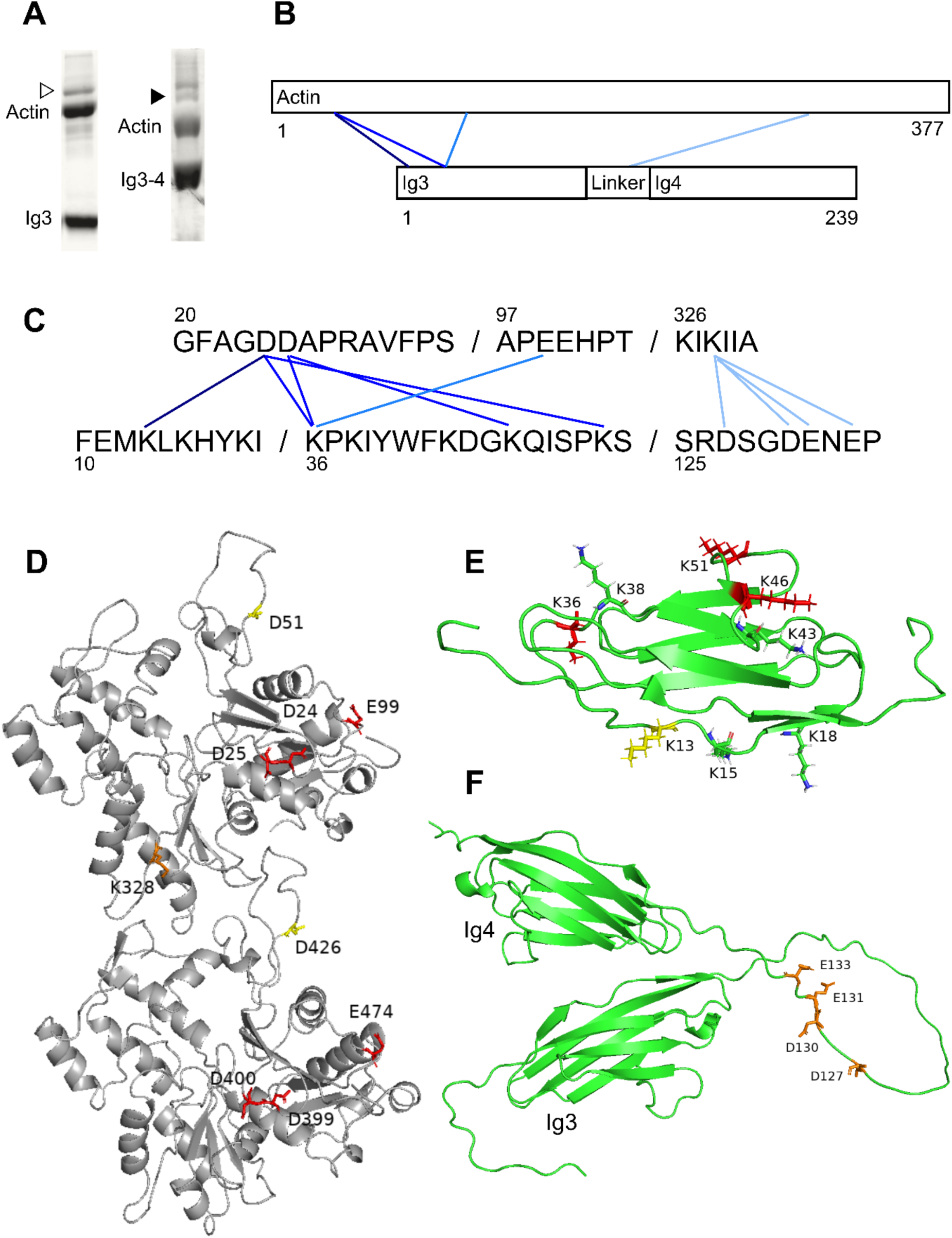
Overview of crosslinks and residues involved in palladin to actin binding as determined by XL-MS. (A) Representative SDS-PAGE gel lanes showing 1:1 palladin:actin crosslinked species. Open triangle indicates Ig3WT (12 kDa) and actin (42 kDa) oligomer at 54 kDa. Closed triangle indicates Ig3-4WT (26.4 kDa) and actin oligomer at 68 kDa. (B) Interprotein crosslinks shown as blue lines indicating general positioning and (C) enlarged to show specific linked residues between actin (top row) and Ig3-4 (bottom row) with lines indicating specific links. (D) Actin dimer and (E) Ig3 showing positions of residues involved in the binding interface. Red residues were included in Pool 1 for HADDOCK trials and yellow residues were also included in Pool 2. Surface exposed lysine residues on Ig3 not found in this study are green. Orange residues were found in crosslinks with the Ig3-4 linker region. (F) Orientation of palladin Ig3-4 as predicted by AlphaFold. Labeled residues in the linker were identified in crosslinks by XL-MS.

**Table 1:** Inter-protein links between actin and the palladin constructs as identified by chemical crosslinking followed by LC-MS/MS and pLink processing. Number in parentheses indicates how many trials per construct identified the link. Links identified by fewer than three total trials are omitted. **Crosslinks included in Pool 1. *Crosslinks included in Pool 2.

| Actin residue | Palladin residue | Palladin construct |
| --- | --- | --- |
| D24** | K36 | Ig3 (3/3), Ig3-4 (4/4) |
| D24** | K51 | Ig3 (3/3), Ig3-4 (4/4) |
| D25** | K36 | Ig3 (3/3), Ig3-4 (4/4) |
| D25** | K46 | Ig3 (2/3), Ig3-4 (4/4) |
| E99** | K36 | Ig3 (3/3), Ig3-4 (4/4) |
| D24* | K13 | Ig3 (2/3), Ig3-4 (2/4) |
| D51* | K46 | Ig3 (1/3), Ig3-4 (3/4) |
| E99 | K46 | Ig3 (2/3), Ig3-4 (1/4) |
| E100 | K36 | Ig3 (2/3), Ig3-4 (1/4) |
| K328 | E133 | Ig3-4 (4/4) |
| K328 | E131 | Ig3-4 (4/4) |
| K328 | D130 | Ig3-4 (4/4) |
| K328 | D127 | Ig3-4 (4/4) |

**Table 2:**
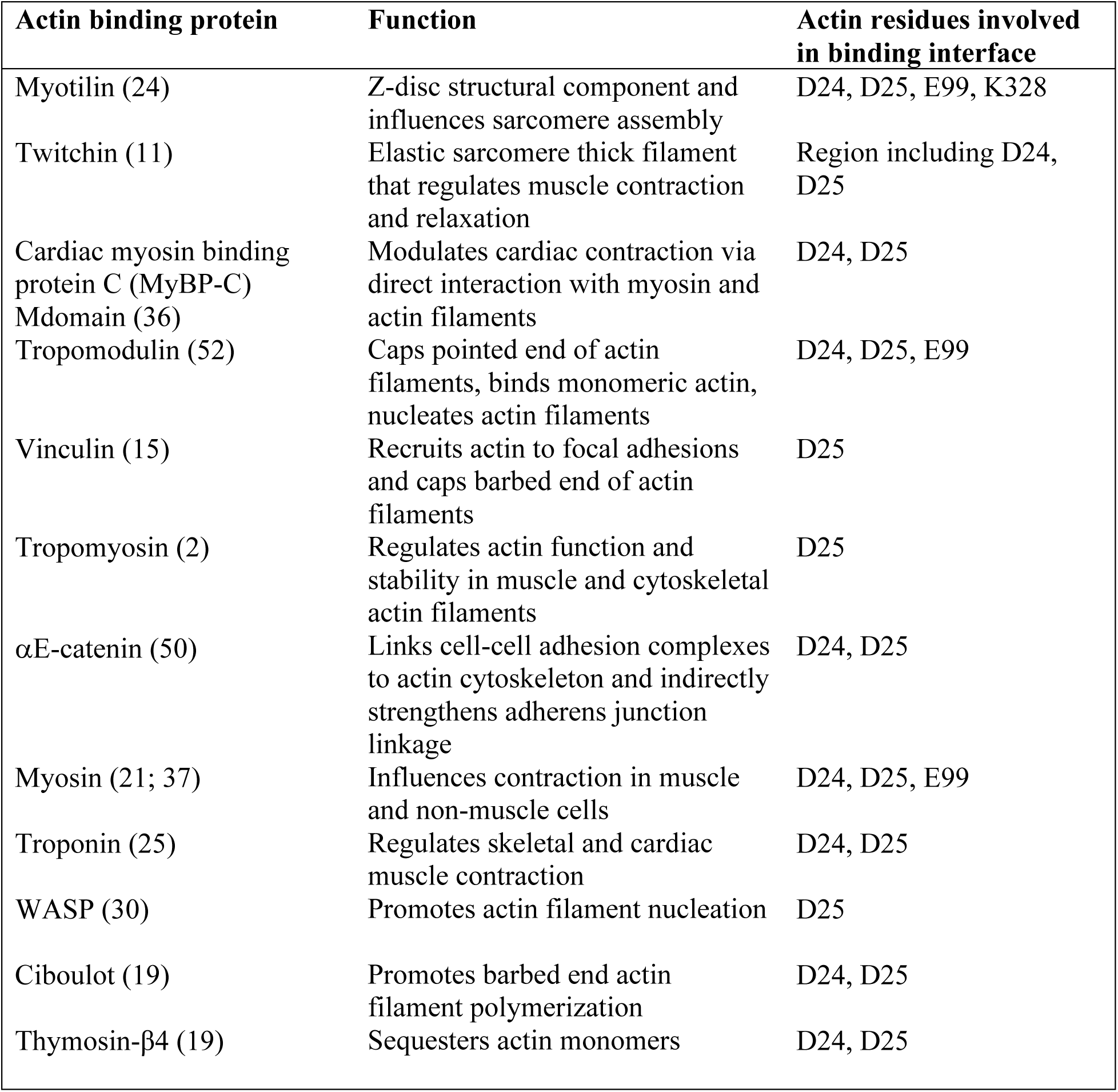
List of actin binding proteins that include shared actin residues in the binding interface.

| **Actin binding protein** | **Function** | **Actin residues involved in binding interface** |
| --- | --- | --- |
| Myotilin (24) | Z-disc structural component and influences sarcomere assembly | D24, D25, E99, K328 |
| Twitchin (11) | Elastic sarcomere thick filament that regulates muscle contraction and relaxation | Region including D24, D25 |
| Cardiac myosin binding protein C (MyBP-C) Mdomain (36) | Modulates cardiac contraction via direct interaction with myosin and actin filaments | D24, D25 |
| Tropomodulin (52) | Caps pointed end of actin filaments, binds monomeric actin, nucleates actin filaments | D24, D25, E99 |
| Vinculin (15) | Recruits actin to focal adhesions and caps barbed end of actin filaments | D25 |
| Tropomyosin (2) | Regulates actin function and stability in muscle and cytoskeletal actin filaments | D25 |
| αE-catenin (50) | Links cell-cell adhesion complexes to actin cytoskeleton and indirectly strengthens adherens junction linkage | D24, D25 |
| Myosin (21; 37) | Influences contraction in muscle and non-muscle cells | D24, D25, E99 |
| Troponin (25) | Regulates skeletal and cardiac muscle contraction | D24, D25 |
| WASP (30) | Promotes actin filament nucleation | D25 |
| Ciboulot (19) | Promotes barbed end actin filament polymerization | D24, D25 |
| Thymosin-β4 (19) | Sequesters actin monomers | D24, D25 |

Prior to use of the identified crosslinks, HADDOCK 2.4 (47) was used to dock Ig3 to F-actin with no restraints. To utilize these crosslinks between Ig3 and F-actin for integrative structural modeling, consistent links were divided into two pools (Table 1). Pool 1 includes all links seen in at least six of seven samples, while pool 2 includes links found in at least four of seven samples. Links found in less than four samples were not included in any docking experiments. Docking of actin side-binding proteins is further complicated by the inability to determine which actin subunit the crosslinks are bound to in the polymer. Therefore, actin subunits were labeled with sequential residue numbering, and all crosslinks were duplicated before macromolecular docking using DisVis (45; 46; 20) and HADDOCK 2.4. DisVis was first utilized to filter out false positives and Xwalk scores were calculated for all HADDOCK-generated structures to validate “crosslinkability” as described (23). All the top scoring structures positioned the Ig3 domain in the cleft between the actin monomers; however, the exact orientation of the domain varied slightly as shown in Figures 2A-C which show the orientations with the best HADDOCK and Xwalk scores, respectively. We also evaluated the top-scoring models from HADDOCK when no XL-MS restraints were used as shown in Supplemental Figure S1 and Table S1. Both the docked structure without restraints and the top scoring structures when the crosslinks were used showed Ig3 in the same cleft between actin monomers. A second round of docking was performed after removing the links with a solvent accessible surface distance (SASD) greater than 50 Å within each HADDOCK cluster as determined by Xwalk and repeating the docking with the original parameters (Figure 2D-F). However, the crosslinks identified and removed by the Xwalk scores were not consistent between the HADDOCK runs and there was no significant correlation between HADDOCK and Xwalk score (Table S1). Yet, the second round of docking output structures had more varied conformations with no clear, significant increase in HADDOCK or Xwalk scores (Table S2). In the second round of docking, when the restraints were defined as ambiguous, noticeably higher HADDOCK scores and lower Xwalk scores were observed. The reverse trend was observed where restraints were classified as unambiguous. Overall, the initial round of docked structures produced from the ambiguous Pool 1 restraints yielded the best overall combination of HADDOCK and Xwalk scores.

**Figure 2:**
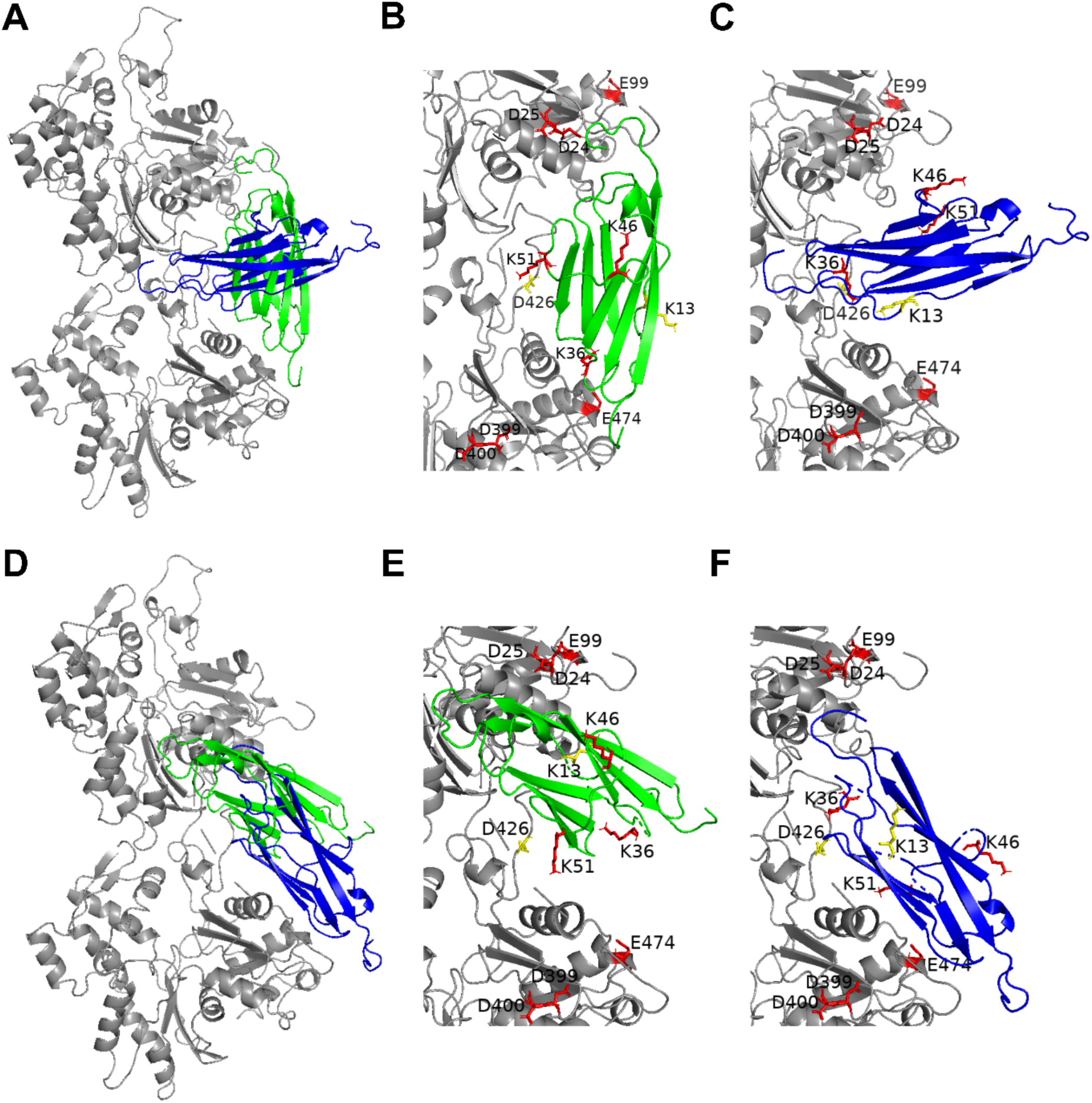
Top scoring models from HADDOCK trials based on HADDOCK and Xwalk score. Blue structures have the highest Xwalk score and green structures have the highest HADDOCK score. (A) Best models from initial trials showing Ig3 positioning between two actin monomers and (B, C) enlarged view with labeled residues. (D) Best models from refined trials showing Ig 3 positioning between two actin monomers and (E, F) enlarged view with labeled residues. For all structures, red residues were included in Pool 1 for HADDOCK trials and yellow residues were also included in Pool 2.

### Validation of lysine residue involvement in actin binding

While prior research has indicated two basic patches on opposite faces of Ig3 are involved in actin binding (3), this new structural model of the complex between palladin’s Ig3 domain and actin indicates additional basic residues are involved in this interaction. We set out to test these predictions by generating point mutations at K13, K36, and K46, which were each consistently identified in crosslinks with actin and had not previously been shown to be involved in F-actin binding. Mutations at K13 and K36 significantly reduced the affinity for F-actin with respect to the WT Ig3 domain (Figure 3). Collectively, these results validate the structural model and corroborate the involvement of electrostatic interactions involving multiple basic patches on palladin with acidic surface of actin.

**Figure 3:**
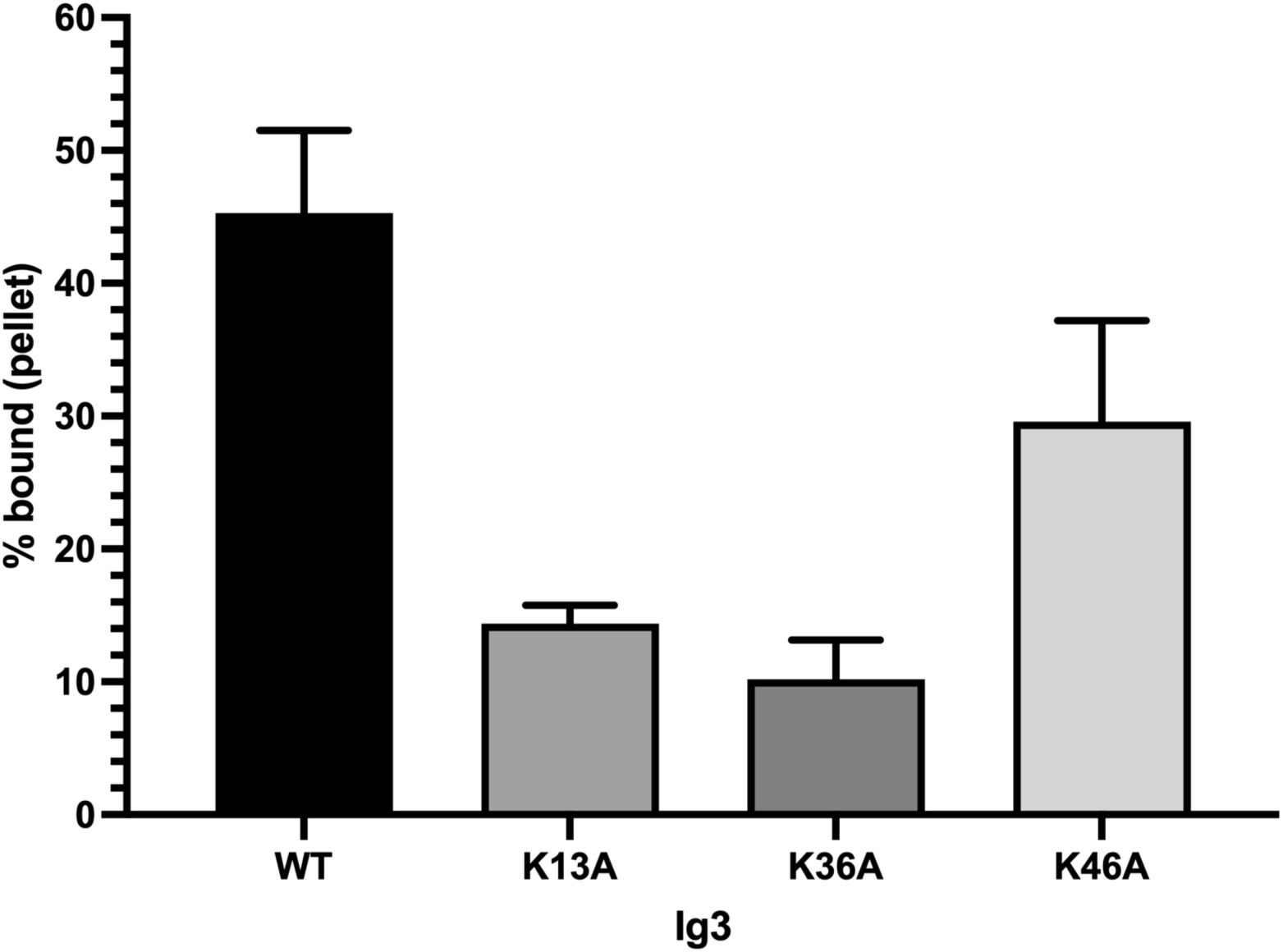
Binding of WT and mutant forms of Ig3 to F-actin. Graph data are presented as a percentage of total binding corresponding to the total amount of protein (10 μM Ig3) with 20 μM F-actin. Data represent mean values ± SEM of 3 independent experiments. The mean binding to F-actin of all mutants except K46A were significantly different from the means of WT Ig3. Significance was assessed using 1-way ANOVA with Holm-Sidak test, *p*<0.001 for WT vs. all other mutants.

**Table 3:** Comparison of molecular weight and R_max_ values from SAXS for palladin constructs.

| Protein | M.W. (kDa) | $R_{\max}$ (Å) |
| --- | --- | --- |
| Ig3 | 11.9 | 67 |
| Ig4 | 11.7 | 68 |
| Ig3-4 | 26.5 | 123 |

### Ig3 and Ig4 domains display conformational plasticity

Both AlphaFold and Rosetta predictions of the structure of the tandem Ig3-4 domain suggest that the linker region is highly disordered (Figure 1F); however, it is unclear how the two Ig domains orient with respect to one another. To investigate interdomain dynamics and flexibility of Ig3-4, SAXS data was collected for the tandem domains as well as each isolated domain in solution. We monitored the maximum scattering intensities extrapolated to zero angle (I_o_) for a concentration series of Ig3, Ig4, and Ig3-4 and observed a minimal concentration dependence for each construct (Figure S2). Therefore, two data curves were merged per protein as described in the methods section. Analysis of the Guinier region for each protein showed a linear fit with residuals distributed evenly around zero indicating monodisperse samples without interparticle interactions (Figure S3). The normalized Kratky plots provide information on the compactness and flexibility of each protein (Figure 5A). All proteins have a peak around 0.1-0.15 Å^−1^ that is indicative of a globular protein. Each data curve also dips around 0.3 Å^−1^ but does not return to baseline which implies flexibility. The pair distance distribution function (PDDF) plot shows single peaks for Ig3 and Ig4 indicating a spherical shape or globular protein as expected (Figure 5B). The Ig3-4 PDDF shows one main peak as well as a shoulder peak around 40-45 Å which suggests a dumbbell shape that is typical for a multidomain protein with two relatively isolated domains connected by a flexible linker (Figure 5B).

**Figure 4:**
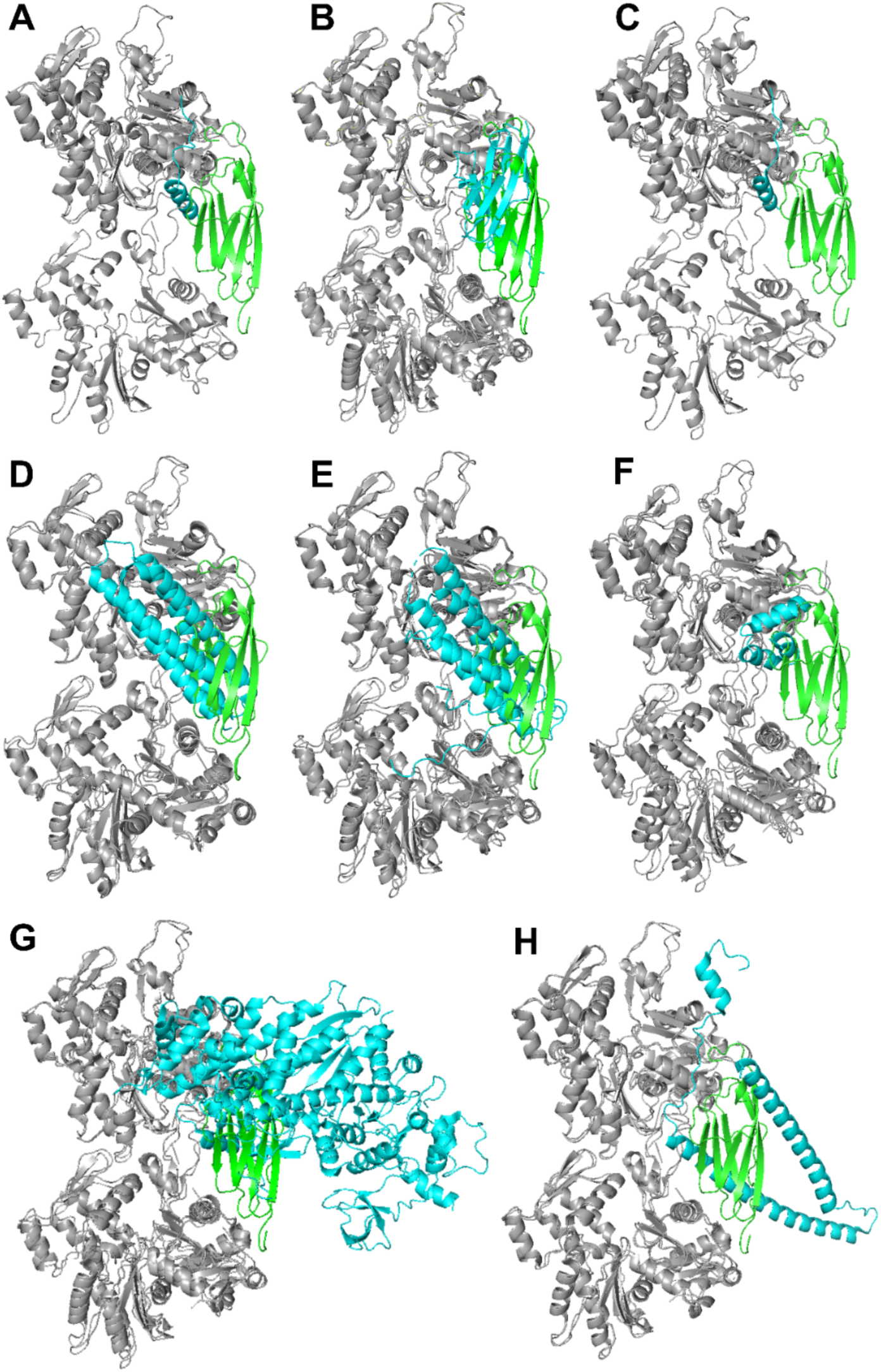
Overlay of docked Ig3 (green) to actin (grey) structure with other ABPs that bind to the same actin residues (cyan). (A) ciboulot, PDB: 1SQK; (B) MyBP-C C2 domain, PDB: 7LRG; (C) N-WASP VC domain, PDB: 2VCP; (D) vinculin tail domain, PDB: 3JBI; (E) αE-catenin, PDB: 6WVT; (F) MyBP-C M-domain, PDB: 7TIJ; (G) myosin, PDB: 7JH7; and (H) troponin C, PDB: 8UZX.

**Figure 5:**
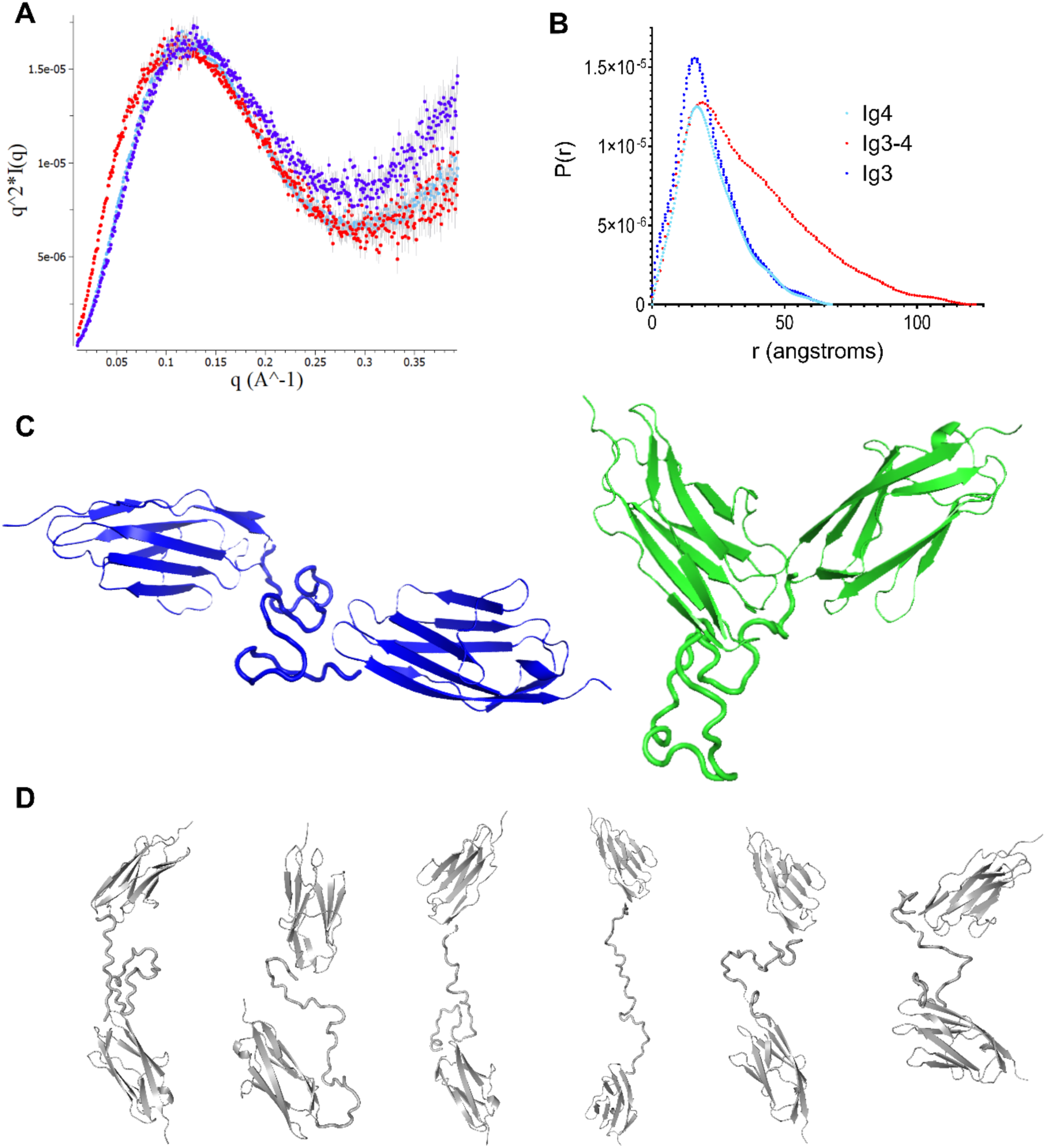
SAXS structural data and EOM structures. (A) Kratky plot and (B) PDDF of Ig3 (blue), Ig4 (cyan), and Ig3-4 (red). (C) The two most likely conformations of Ig3-4 as predicted by EOM with each contributing approximately 25% fractional occupancy. (D) The remaining six conformations of Ig3-4 as predicted by EOM with each contributing approximately 8% fractional occupancy.

Data quality was further assessed by using the FoXS online web server to compare experimental data to theoretical data generated from the solved NMR structures for each of the single domain samples. Scattering curves from Ig3 and Ig4 showed high similarity with χ^2^ values of 3.17 and 10.6, respectively (Figure S4A-B). The structure of Ig3-4 was excised from the predicted palladin structure from AlphaFold (Q9ET54) and compared to the collected Ig3-4 data (Figure S4C). A much higher χ^2^ value of 38.9 suggests this predicted orientation does not agree well with the experimental data collected in this study. Similarly, comparison with CRYSOL between the experimental data and AlphaFold structure produces a high χ^2^ value of 85.471. SASREF quaternary structural modeling run using CRYSOL amplitude files from the individual domains versus the Ig3-4 experimental scattering curve also shows low correlation between the data and AlphaFold structure.

To account for the likely flexibility between Ig3 and Ig4 within the tandem domain, Ensemble Optimization Method (EOM) was performed. Sub-programs RANCH and GAJOE created an ensemble of possible conformations, and a genetic algorithm determined the conformations consistent with the experimental SAXS data. The best fit to the data produced a χ^2^ value of 1.077, and the fit between experimental and theoretical scattering as well as distribution plots for D_max_, N-C distance, and Rg distance are included in Supplemental Figure S5. Two conformations contributed equally to the final ensemble each with approximately 25% fractional occupancy. The first of these structures presents the Ig domains in a “V” orientation with the unstructured linker region extending from the bottom of the “V”. The second orientation shows a linear orientation between the domains with the linker region between the domains (Figure 5C). The remaining six structures in the ensemble each compose approximately 8% fractional occupancy with intermediate conformations (Figure 5D).

### Ig3 and Ig4 domain residues interact within the tandem Ig3-4 construct

One factor that could enhance the F-actin binding affinity and bundling efficiency of Ig3-4 versus the minimal actin binding domain could be an interaction between the Ig3 and Ig4 domains. To identify potential interactions, we collected separate ^1^H-^15^N HSQC spectra of Ig3, Ig4, and Ig3-4 (Figure 6A) and calculated the chemical shift perturbation (CSP) of residues from the individual versus tandem domains (Figure 6B). CSP values above a cutoff of 0.05 ppm were considered significant. While high CSP values near the C-terminus and central linker region were not considered due to the likely flexibility of these regions, the chemical shifts of several residues within both the Ig3 and Ig4 regions significantly differed between the individual domains and tandem construct as highlighted in Figure 6C and D. Much more significant chemical shift perturbations were observed in Ig3 domain residues in comparison with Ig4 domain. This result is consistent with our previous result in which Ig3-4 forms an intramolecular crosslinked species (48). This data also identifies the potential involvement of the following residues in the interdomain interactions between Ig3 and Ig4 as: G22, T26, H54, T56, H68, T70, T73, D76, L97, F144, L168, H198, A221, and F226. An additional observation is that neither the lipid-binding nor actin-binding regions of the Ig3 domain overlap with the Ig4 domain interacting regions (51). This suggests that the interaction between the Ig3 and Ig4 domains would not interfere with lipid or actin binding.

**Figure 6:**
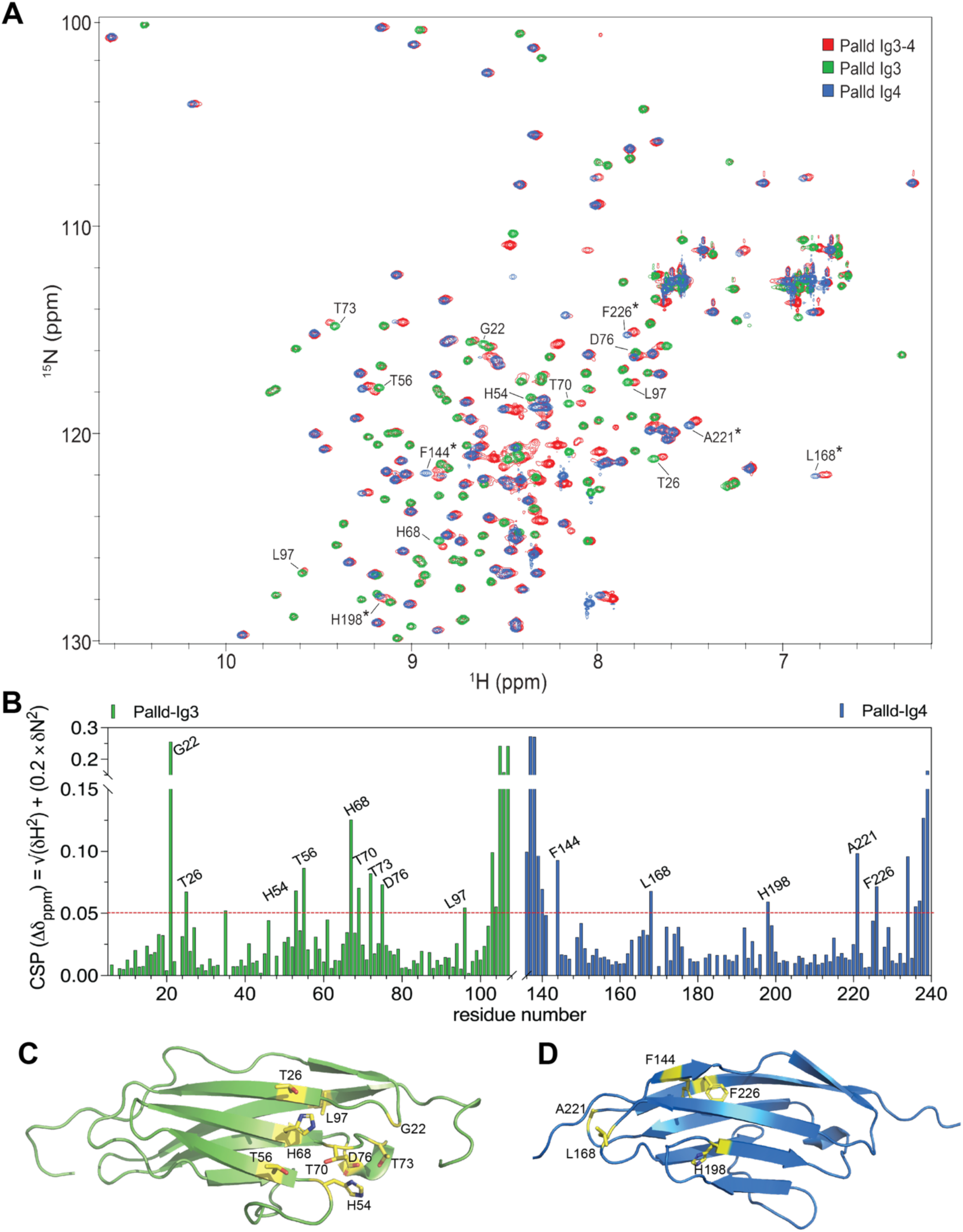
NMR results showing interaction between Ig3 and Ig4 domain residues. (A) HSQC spectra overlay of Ig3-4 (red), Ig3 (green), and Ig4 (blue). Labeled residues were found to have a significant CSP between Ig3-4 and Ig3 (no *) or Ig4 (*). (B) Chemical shift perturbation of residues from comparison of Ig3-4 with the individual domains. Red dotted line indicates cutoff value over which CSPs were deemed significant. Residues above the cutoff value are highlighted in the (C) Ig3 and (D) Ig4 domains.

## Discussion

Here, we have defined the molecular basis through which palladin binds to the sides of actin filaments (Figures 2 and 4A). A combination of XL-MS, NMR, SAXS, and mutagenesis studies indicate how distinct basic patches within the Ig3 domain of palladin contribute to bind actin. Previous work had established that the minimal domain required for F-actin binding by palladin is the Ig3 domain (9). Several lysine residues within two basic patches on the surface of palladin’s Ig3 domain have been identified as being critical for F-actin interactions. Initially, mutagenesis studies identified K15 and K18 as one basic patch and K51 on the opposite face of Ig3 to be crucial for F-actin binding (3). However, mutagenesis of individual residues was not enough to eliminate all F-actin binding which required double and triple mutants that also resulted in nuclear localization when expressed in cells. A later study found that K38, which is near K51 in the structure, may be part of the second basic patch with K51 and influence actin polymerization and bundling (51). Here, mass spectrometry identification of chemically crosslinked peptides of the Ig3:actin complex revealed that K13, K36, K46, and K51 are in close proximity to D24, D25 and E99, all residues of actin’s most negatively charged subdomain 1 (44). Reduced actin binding by all four single point mutations at these lysine residues substantiate that there are two different basic patches that interact at the interface between two actin monomers in an orientation like several other side-binding proteins (Figure 4). This structure has implications for the multiple functions and regulation of actin binding, bundling and polymerization by palladin.

Protein regulation works by adjusting the conformational states which are often regulated by binding interactions. Our previous study of Ig3 and Ig3-4 with membrane phosphoinositide, PI(4,5)P_2_, indicated that the binding of PI(4,5)P_2_ reduces the actin-binding and polymerizing activity of both Ig3 and Ig3-4 (51). Moreover, the chemical shift perturbation experiment of Ig3 and inositol 1,4,5-trisphosphate (IP_3_) pinpointed a region involving residues K38, Q47, and K51 in this interaction (51). The similarity of the region involved in both interactions (Ig3/actin and Ig3/PI4,5P_2_) and the effect of PI4,5P_2_ binding in reduction of Ig3/Ig3-4 actin binding and polymerizing activity reinforces an essential role of the region involving K36, K46, and K51 in actin binding and its activity.

We also showed that the chemical shift perturbations observed for the Ig3-4 tandem domain, when compared with the isolated Ig3 and Ig4 domains, identified a region (residues T26, H54, T56, H68, T70, and T73) that does not overlap with the Ig3:actin interacting residues identified in this manuscript or the Ig3:lipid interacting residues previously identified (51). This suggests that the interaction between the Ig3 and Ig4 domains would not interfere with lipid or actin binding. Furthermore, the ensemble Ig3-4 PDB model based on SAXS data shown in Figure 5C (green) substantiates that the interface between the Ig3 and Ig4 domains does not overlap with Ig3’s actin-binding site. In addition, it is also likely that Ig3-4 could adopt a more extended conformation in the presence of actin or that fractional occupancy of other conformations could change upon actin binding favoring higher actin binding and bundling activity by Ig3-4 in comparison to Ig3 alone.

Palladin and actin binding dynamics are further complicated by the formation of a palladin homodimer in the presence of actin, and prior chemical crosslinking studies found that palladin Ig3-4 tandem domain forms an intramolecular crosslinked species even in the absence of actin (48). These intramolecularly crosslinked species showed reduced ability to bind and bundle actin. Cleavage of the crosslinking agent used partially restored binding and bundling ability despite remaining modifications on the lysine residues indicating that flexibility and orientation between Ig3 and Ig4 is critical for these functions. In this study, we identified several positions in the linker (D127, D130, E131, and E133) that are involved in crosslinks to actin at K328. While these residues may not be specifically involved in interactions between Ig3 and Ig4, the fact that they were able to be crosslinked indicates that some surface exposed residues could be obstructed in this intramolecularly linked species. This also raises the possibility that binding to actin induces a conformational change within Ig3-4 exposing a dimerization site. Further research into the specific mechanism of palladin dimerization and interactions between Ig3 and Ig4 is clearly needed.

While Ig3 is the only palladin domain known to bind to actin, the enhanced binding and bundling ability of Ig3-4 warrants further investigation. The mechanism for this amplification could be due to either the presence of the Ig4 domain or of the unique, unstructured linker region. Our data presented here suggests that both of these factors contribute. The XL-MS data provides the first evidence of an interaction between the residues in palladin’s linker and residues on the surface of actin. HSQC spectroscopy shows CSPs of residues in both the Ig3 and Ig4 domains indicating potential binding between the domains. It has previously been found that both Ig3 and Ig3-4 form homodimers in the presence of actin as well as an intramolecularly crosslinked Ig 3-4 species. XL-MS of these dimeric and crosslinked species would provide further insight into potential binding between palladin’s Ig domains. Little research has been conducted on the linker region, but several factors could be of impact including its extended length, overall charge, and specific composition of amino acids.

Various lines of evidence support the notion that basic residues on the surface of the Ig3 domain of palladin are critical for actin binding, displaying a binding mode similar to several other side-binding proteins as shown in Figure 4. Yet, the functions of these actin-binding proteins vary widely, as highlighted in Table 2. Palladin, as the founding member of a unique family of actin-binding proteins that are involved in controlling the architecture and dynamics of diverse and versatile actin-based structures, displays a wide variety of roles. Besides binding and crosslinking actin filaments, palladin has also been shown to promote nucleation of actin polymerization (18) and can functionally replace the Arp2/3 complex in *Listeria* actin comet tail motility (8). Based on our results, we present a structural model that is consistent these diverse roles (Figure 7). In conclusion, our results clearly show that the Ig3-4 domain pair of palladin is flexible, given the unusually long linker region, and is thus able to accommodate both an extended, linear and a V-shaped conformation with possible implications for different functions in the cell as indicated in our model (Figure 7). Although both the actin-binding interface and Ig domain dynamics are similar to what was observed for myotilin (34; 24), further studies are required to determine how this family of proteins carry out both overlapping and distinct functions in a wide array of cellular roles.

**Figure 7:**
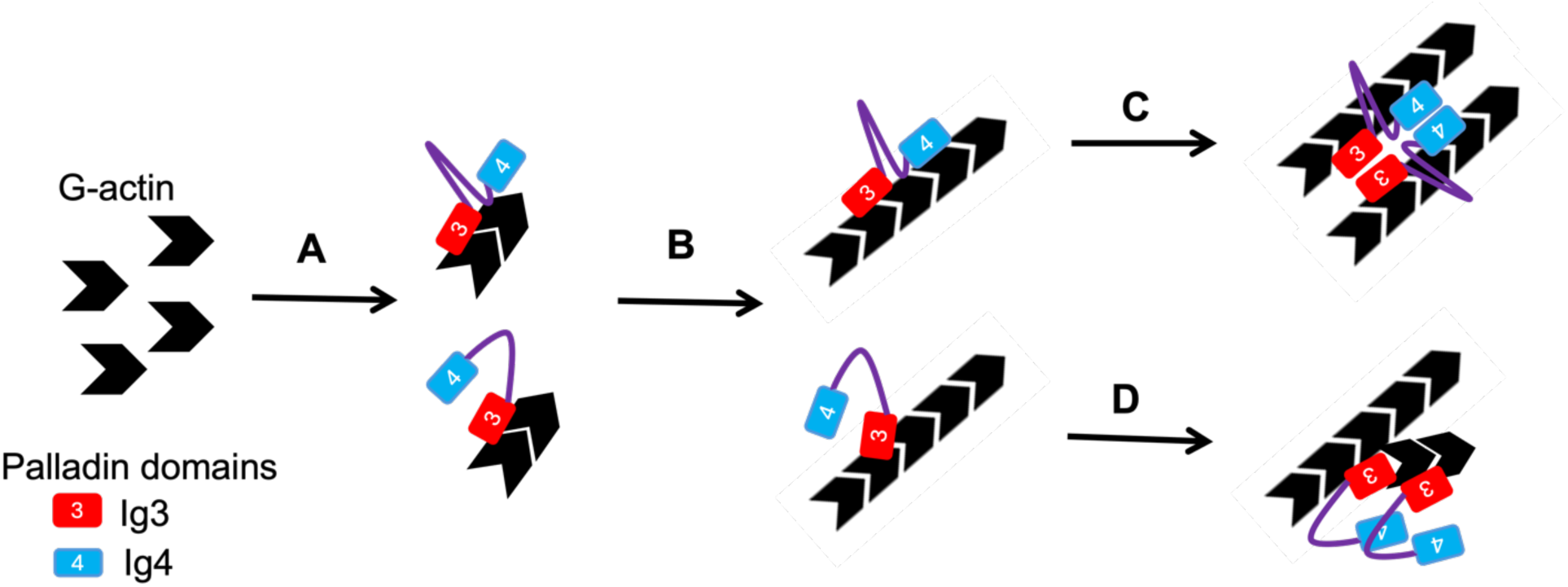
Model illustrating multiple roles of palladin in actin dynamics and organization. (A) Palladin Ig3 domain binds and stabilizes actin seeds to promote nucleation step of polymerization by binding between actin protomers. (B) A conformational change occurs upon binding actin, that promotes dimerization of palladin that can result in (C) actin bundling or (D) branching.

## Materials and Methods

### Protein Expression and Purification

Palladin domains were expressed and purified from BL21(DE3) *Escherichia coli* cells as previously described (18). Several lysine residue mutants (K13A, K36A, K46A, and K51A) were generated from the pTBSG WT Ig3 construct using the Q5^®^ site-directed mutagenesis kit (New England Biolabs). Sequences were verified by DNA sequencing (Supplemental Note 1 provides each protein sequence). Briefly, the WT and mutant Ig3 and Ig4 domains in the pTBSG expression vector were expressed in *E. coli* using autoinduction media (43) and were subsequently lysed by sonication and centrifuged to pellet cell debris. Supernatant containing soluble protein was purified using HisPur Ni-NTA resin (Thermo Scientific). The His6-tag was cleaved using tobacco etch virus (TEV) protease and further purified by cation exchange chromatography (SP Sepharose, GE Healthcare Life Sciences). Protein was dialyzed to HEPES storage buffer (20 mM HEPES, pH 7.4, 1 mM DTT, 100 mM NaCl). The Ig3-4 tandem domain was expressed from the pTBMalE expression vector that in addition to the His6-tag contains a maltose binding protein (MBP) tag for increased solubility. Ig3-4 was purified using the same method as Ig3/Ig4 with the addition of an amylose column (New England Biolabs) chromatography step to remove the MBP tag prior to cation exchange. Actin was purified from rabbit muscle acetone powder (Pel-Freez Biologicals) using an adapted method from Spudich and Watt (42).

### Chemical Crosslinking

Actin was polymerized in 2X polymerization buffer (20 mM Tris at pH 8.0, 200 mM KCl, 2 mM MgCl_2_, and 4 mM DTT) for at least 30 minutes at room temperature. F-actin (final concentration 10 μM) was incubated with varying amounts of DMTMM (2 or 5 mM) for 5 minutes at room temperature. Purified palladin (10-20 μM) in HEPES storage buffer was added, and the mixture was incubated for a further 40 minutes at room temperature. Next, 4X Laemmli buffer (200 mM Tris-HCl, pH 6.8, 8% SDS, 40% glycerol, 0.08% bromophenol blue, and 400 mM DTT) was added to quench the reaction and samples were boiled and separated by SDS-PAGE. Protein bands at a molecular weight according to a 1:1 ratio of palladin to actin were excised from the gel.

### Gel-Based MS/MS Analysis

Each SDS-PAGE gel band was subjected to in-gel trypsin digestion as follows. Gel segments were destained in 50% methanol (Fisher), 50 mM ammonium bicarbonate (Sigma-Aldrich), followed by reduction in 10 mM Tris[2-carboxyethyl]phosphine (Pierce) and alkylation in 50 mM iodoacetamide (Sigma-Aldrich). Gel slices were then dehydrated in acetonitrile (Fisher), followed by addition of 100 ng porcine sequencing grade modified trypsin (Promega) in 50 mM ammonium bicarbonate (Sigma-Aldrich) and incubation at 37 °C for 12-16 hours. Peptide products were then acidified in 0.1% formic acid (Pierce). Tryptic peptides were then separated by reverse phase XSelect CSH C18 2.5 um resin (Waters) on an in-line 150 x 0.075 mm column using an UltiMate 3000 RSLCnano system (Thermo). Peptides were eluted using a 75 min gradient from 98:2 to 65:35 buffer A:B ratio where Buffer A is 0.1% formic acid and 0.5% acetonitrile and Buffer B is 0.1% formic acid and 99.9% acetonitrile. Eluted peptides were ionized by electrospray (2.4 kV) followed by mass spectrometric analysis on an Orbitrap Eclipse Tribrid mass spectrometer (Thermo). MS data were acquired using the FTMS analyzer in profile mode at a resolution of 120,000 over a range of 375 to 1400 m/z with advanced peak determination. Following HCD activation, MS/MS data were acquired using the FTMS analyzer in profile mode at a resolution of 15,000 over a range of 150 to 2000 m/z with a stepped collision energy of 27-33%.

### Analysis and visualization of LC-MS/MS data

pLink 2 version 2.3.11 software (5) was used for identification of crosslinked peptides with searches performed against the forward and reverse protein sequences. Raw data files were analyzed using the default options with the following exceptions: EDC-DE was used as the crosslinker, Carbamidomethyl[C] was fixed, Oxidation[M] was variable, and E-values were computed. Results were filtered at a tolerance of ± 10 ppm and a false discovery rate (FDR) of 5% at peptide spectrum matches (PSM) level. xiNET cross-link viewer (6) was used for visualization of the crosslinks identified by pLink2. Results were uploaded to the Proxl (Protein Cross-Linking Database) server (35) and link to data is provided in Supplemental Note 2.

### Protein docking with DisVis and HADDOCK

The interprotein crosslinks identified by pLink 2 were sorted into three categories: crosslinks present in at least six of seven trials (Pool 1), crosslinks present in at least four of seven trials (Pool 2), and crosslinks present in less than four trials. Crosslinks present in less than four trials were not utilized. For docking, all steps were performed separately for both Pool 1 and Pool 2. Because palladin is hypothesized to bind between two actin monomers within a filament and because there is no way to differentiate which monomer was actually linked, all crosslinks were duplicated to the second monomer. For actin, a dimer of chains D and F was extracted from PDB 3J8I and reformatted as one chain with sequential residue numbering. The first structure of the palladin Ig3 structural ensemble PDB 2LQR was used for all modeling. Links were reformatted as required by the docking software indicating the PDB file chain ID, residue number of the crosslink, atom position of the crosslink, and upper and lower distance restraints (23 and 24 Å for Lys-Asp and Lys-Glu links, respectively). Distance restraints were calculated by summing the Euclidian distance between the α-carbon and the most distant atom on each residue in each crosslink with 13 Å added for conformation changes (29). DisVis software was used to identify the accessible interaction space and to detect crosslinks that violate the distance restraints. Crosslinks with a DisVis Z-score of less than 0.5 were used in HADDOCK (47; 20) and trials were run in duplicate assigning crosslinks as either ambiguous or unambiguous. Center of mass restraints were selected. Best models were identified with help from the workflow described by Orbán-Németh *et al.* (29). Xwalk (23) was used on the resulting structures to calculate the Solvent Accessible Surface Distance (SASD) using only backbone and beta carbon atom coordinates, solvent radius set to 2, and a maximum distance of 50 Å with only inter-molecular distances output. Restraints were identical to HADDOCK runs. Further refinement was accomplished by removing crosslinks with a SASD greater than 28 Å within each HADDOCK cluster as determined by Xwalk and docking again in HADDOCK using the original parameters. Resulting structures were then scored again by Xwalk.

### Actin co-sedimentation assays

Actin binding properties of the three Ig3 variants (K13A, K36A, and K46A) were compared to WT using an actin co-sedimentation assay (9). Briefly, we incubated 10 μM Ig3 protein with 20 μM F-actin that were centrifuged at high speed (150,000 xg) for 30 min before separating the supernatant and pellet. Actin and Ig3 proteins in the supernatant and solubilized pellet were separated using 12% SDS-PAGE and protein bands were quantified using Fiji software (39). All binding assays were repeated at least three times and averaged data with standard deviation error bars are presented.

### Small Angle X-ray Scattering

After dialysis of purified palladin domains to HEPES storage buffer, proteins were further purified by size exclusion chromatography on the Cytiva Superdex 200 10/300 GL column in preparation for SAXS data collection. Ig3 and Ig4 were concentrated and prepared directly for SAXS, while Ig3-4 was subjected to a final purification step on the Cytiva HiLoad 16/600 Superdex 200 pg column to remove the remaining contaminants. Concentration series were prepared for each protein: 0.5-4.3 mg/mL for Ig3, 0.5-10 mg/mL for Ig4, and 0.5-7.4 mg/mL for Ig3-4. SAXS data was collected at twelve frames per sample with 1 second exposures at the Stanford Synchrotron Radiation Library (SSRL) (41). Resulting data was automatically buffer-subtracted and analyzed on-line with SAXSPipe. The ATSAS software package was used for all further analysis (10; 27). Buffer-subtracted files were compared at every concentration for each protein. Slight concentration dependence was observed for all domains, so the lower q range of a lower concentration sample was merged with the higher q range of a higher concentration sample using PRIMUS. PRIMUS was further used for each merged data curve to estimate radius of gyration (Rg) and the forward scattering intensity I(0) from the Guinier plot and Porod volume and Rmax from the distance distribution analysis.

Prior to structural modeling, sequence and structure files were optimized as follows. For Ig3, the N-terminal “SNA” and “GGS” residues were removed from the sequence and pdb files, respectively, for sequence continuity. For Ig4, the C-terminal “AHK” and “SGPSSG” residues were removed from the sequence and pdb files, respectively. Furthermore, the sequence was modified with I179V and N224S in order to match the structure.

To check data quality for Ig3 and Ig4 with known structures, CRYSOL (maximum *s* value 0.4), SASREF, and the FoXS webserver were used to create and compare theoretical scattering curves to the experimental data and to create an initial rigid-body model. The AlphaFold predicted structure for Ig3-4 was used for experimental data comparison with FoXS. SASREF was run with the Ig3-4 experimental data and amplitudes for the individual domains were computed by CRYSOL. EOM was employed by using the sub-program RANCH to generate a pool of 10,000 possible structures from sequence and pdb files with a maximum *s* value cut-off of 0.4. Then the sub-program GAJOE was used to compare these models to the experimental data. All programs were run in user mode with default settings unless otherwise specified.

### ^1^H-^15^N HSQC

All NMR experiments were performed at 298 K on a Bruker AVANCE 800 MHz spectrometer equipped with a triple resonance cryoprobe. The NMR buffer was 20 mM Hepes (pH 6.5), 100 mM NaCl, 2 mM DTT, 0.01% NaN_3_, and 10% D_2_O for spectrometer lock. NMR were processed using the NMRPipe program (7) and analyzed and visualized using NMRViewJ (22). A ^1^H-^15^N HSQC spectrum was recorded separately for samples containing 0.35 mM ^15^N labeled Ig3, Ig4, and Ig3-4 domains of palladin. Combined chemical shift perturbations (CSPs) were calculated using the equation Δδ = [((ΔδH)^2^ + (ΔδN/5)^2^ /2]^|1/2|^; where ΔδH and ΔδN are chemical shift changes of ^1^H and ^15^NH, respectively.

## Supplementary Material

SupplementaryData: Supporting information includes Supplemental Note S1 with list of protein sequences, Note S2 with Crosslinking Data Deposition and SASBDB Links, Table S1 with Initial HADDOCK and Xwalk scores, Table S2 with Refined Xwalk and HADDOCK scores, Figure S1 showing HADDOCK structure of complex with no restraints, Figure S2 displaying SAXS concentration series, Figure S3 showing the Guinier region linear fit, Figure S4 showing the fit and distribution plots of RANCH/GAJOE Ig3-4 modeling, and Figure S5 displaying the FoXS scattering curves with X^2^ values.

## Acknowledgments

Research reported in this publication was supported by NIGMS of the National Institutes of Health under award number R15GM120670 and the Kansas INBRE, P20 GM103418. We thank Dr. Haifan Wu (Wichita State University, USA) for assistance with chemical crosslinking knowledge and data analysis software; the UAMS Proteomics Core for XL sample processing and LC/MS-MS; Drs. Thomas Weiss (SSRL, USA) and Allyn Schoeffler (Loyola University New Orleans, USA) for help with SAXS data collection and analysis support; and Dr. Justin Douglas (KU NMR Core, University of Kansas, USA) for assistance with NMR data collection. The FP7 WeNMR (project# 261572), H2020 West-Life (project# 675858), the EOSC-hub (project# 777536) and the EGI-ACE (project# 101017567) European e-Infrastructure projects are acknowledged for the use of their web portals, which make use of the EGI infrastructure with the dedicated support of CESNET-MCC, INFN-LNL-2, NCG-INGRID-PT, TW-NCHC, CESGA, IFCA-LCG2, UA-BITP, TR-FC1-ULAKBIM, CSTCLOUD-EGI, IN2P3-CPPM, CIRMMP, SURFsara and NIKHEF, and the additional support of the national GRID Initiatives of Belgium, France, Italy, Germany, the Netherlands, Poland, Portugal, Spain, UK, Taiwan and the US Open Science Grid. Use of the Stanford Synchrotron Radiation Lightsource, SLAC National Accelerator Laboratory, is supported by the U.S. Department of Energy, Office of Science, Office of Basic Energy Sciences under Contract No. DE-AC02-76SF00515. The SSRL Structural Molecular Biology Program is supported by the DOE Office of Biological and Environmental Research, and by the National Institutes of Health, National Institute of General Medical Sciences (P30GM133894). The contents of this publication are solely the responsibility of the authors and do not necessarily represent the official views of NIGMS or NIH.

## Abbreviations

ABPs: actin binding proteins
Ig: immunoglobulin-like
XL-MS: crosslinking mass spectrometry
SAXS: small angle X-ray scattering
LC: liquid chromatography
PDDF: pair distance distribution function
EOM: Ensemble Optimization Method
NMR: nuclear magnetic resonance
CSP: chemical shift perturbation
PI(4,5)P_2_: phosphatidylinositol 4,5-bisphosphate
SASD: Solvent Accessible Surface Distance

