## Supplemental Data for "Integrated structural model of the palladin-actin complex using XL-MS, docking, NMR, and SAXS"

**Contents**

1 **Supplementary Notes**

Supplementary Note S1. Detailed sequence information of the protein constructs…………....2

Supplementary Note S2. Link for deposition of crosslinking and SAXS data ….………….....3

2 **Supplementary Tables**

Table S1 - Initial HADDOCK and Xwalk scores ……………………………………………..4

Table S2 - Xwalk and HADDOCK scores from refined HADDOCK trials……………….….5

3 **Supplementary Figures**

Figure S1 - HADDOCK structure of complex with no restraints…………………………......6

Figure S2 - SAXS concentration series………………………………………………………..7

Figure S3 - Guinier region linear fit……………………………………………………………8

Figure S4 - FoXS scattering curves with X2 values………………………………...…………9

Figure S5 - Fit and distribution plots of RANCH/GAJOE Ig3-4 modeling…………………..10

**Supplementary Note S1:** Protein Sequences

- **ACTIN** SP|P68135|ACTS_RABIT ACTIN, ALPHA SKELETAL MUSCLE OS=ORYCTOLAGUS CUNICULUS OX=9986 GN=ACTA1 PE=1 SV=1

MCDEDETTALVCDNGSGLVKAGFAGDDAPRAVFPSIVGRPRHQGVMVGMGQKDSYVGDEAQSKRGILTLKYPIEHGIITNWDDMEKIWHHTFYNELRVAPEEHPTLLTEAPLNPKANREKMTQIMFETFNVPAMYVAIQAVLSLYASGRTTGIVLDSGDGVTHNVPIYEGYALPHAIMRLDLAGRDLTDYLMKILTERGYSFVTTAEREIVRDIKEKLCYVALDFENEMATAASSSSLEKSYELPDGQVITIGNERFRCPETLFQPSFIGMESAGIHETTYNSIMKCDIDIRKDLYANNVMSGGTTMYPGIADRMQKEITALAPSTMKIKIIAPPERKYSVWIGGSILASLSTFQQMWITKQEYDEAGPSIVHRKCF

- **Palladin Ig3-4** (26.6 kDa) Mus musculus

SNANATAPFFEMKLKHYKIFEGMPVTFTCRVAGNPKPKIYWFKDGKQISPKSDHYTIQRDLDGTCSLHTTASTLDDDGNYTIMAANPQGRVSCTGRLMVQAVNQRGRSPRSPSGHPHARRPRSRSRDSGDENEPIQERFFRPHFLQAPGDLTVQEGKLCRMDCKVSGLPTPDLSWQLDGKPIRPDSAHKMLVRENGVHSLIIEPVTSRDAGIYTCIATNRAGQNSFNLELVVAAKEAHK

- **Palladin Ig3-4 linker,** Mus musculus

AVNQRGRSPRSPSGHPHARRPRSRSRDSGDENEPIQERFFR

- **Palladin Ig3** (12.0 kDa), Mus musculus

SNANATAPFFEMKLKHYKIFEGMPVTFTCRVAGNPKPKIYWFKDGKQISPKSDHYTIQRDLDGTCSLHTTASTLDDDGNYTIMAANPQGRVSCTGRLMVQAVNQRGRS

Lysines (K13, K36, K46, and K51) mutated to alanine are highlighted in red.

- **Palladin Ig4** (11.8 kDa), Mus musculus

TVQERFFRPHFLQAPGDLTVQEGKLCRMDCKVSGLPTPDLSWQLDGKPIRPDSAHKMLVRENGVHSLIIEPVTSRDAGIYTCIATNRAGQNSFNLELVVAAKEAHK

**Supplementary Note S2:** Data Deposition Links

- Link to Proxl (Protein Cross-Linking Database)

<http://www.yeastrc.org/proxl_public/projectReadProcessCode.do?code=seyd6pmwj5ybwuou17l56tusfalk4v4jowf0or6ni5kveknqjxue9tfgvge8d9rs>

Note: actin residue numbering in the deposited data is shifted +2 compared to the main text

- Link to SASBDB (Small Angle Scattering Biological Data Bank)
  - Draft IDs: 6214 (Ig3), 6213 (Ig3-4), 6212 (Ig4)

**Table S1:** Xwalk and HADDOCK scores from initial HADDOCK trials

| **Crosslinks used** | **Restraint condition** | **HADDOCK cluster** | **HADDOCK score** | **Xwalk score** |
| --- | --- | --- | --- | --- |
| No Restraints | Not applicable | 1 | -52.6 +/- 9.1 | Not applicable |
| Pool 2 | Ambiguous | 1 | -68.6 +/- 6.9 | 0.6173 |
| Pool 1 | Unambiguous | 1 | -75.9 +/- 3.5 | * |
| Pool 2 | Unambiguous | 1 | -11.4 +/- 1.7 | 0.9051 |
| Pool 2 | Unambiguous | 2 | 28.6 +/- 1.7 | 0.9235 |
| Pool 1 | Ambiguous | 1 | -89.0 +/- 5.7 | 0.8643 |

*Could not be determined. No links under 28 Å cutoff.

**Table S2:** Xwalk and HADDOCK scores from refined HADDOCK trials

| **Crosslinks used** | **Restraint condition** | **Initial structure input** | **Best structure output** | **HADDOCK score** | **Xwalk score** |
| --- | --- | --- | --- | --- | --- |
| Pool 2 | Ambiguous | Cluster 2 | Cluster 3_1 | -93.3 +/- 5.8 | 0.8080 |
| Pool 2 | Ambiguous | Cluster 1 | Cluster 5_1 | -92.2 +/- 12.0 | 0.5730 |
| Pool 1 | Unambiguous | Cluster 1 | Cluster 3_1 | -56.6 +/- 10.0 | 0.9259 |
| Pool 1 | Unambiguous | Cluster 1 | Cluster 9_1 | -18.8 +/- 10.3 | 0.9082 |
| Pool 1 | Unambiguous | Cluster 1 | Cluster 6_1 | -18.8 +/- 3.0 | 0.8938 |
| Pool 1 | Ambiguous | Cluster 3 | Cluster 1_1 | -92.7 +/- 12.5 | 0.8732 |

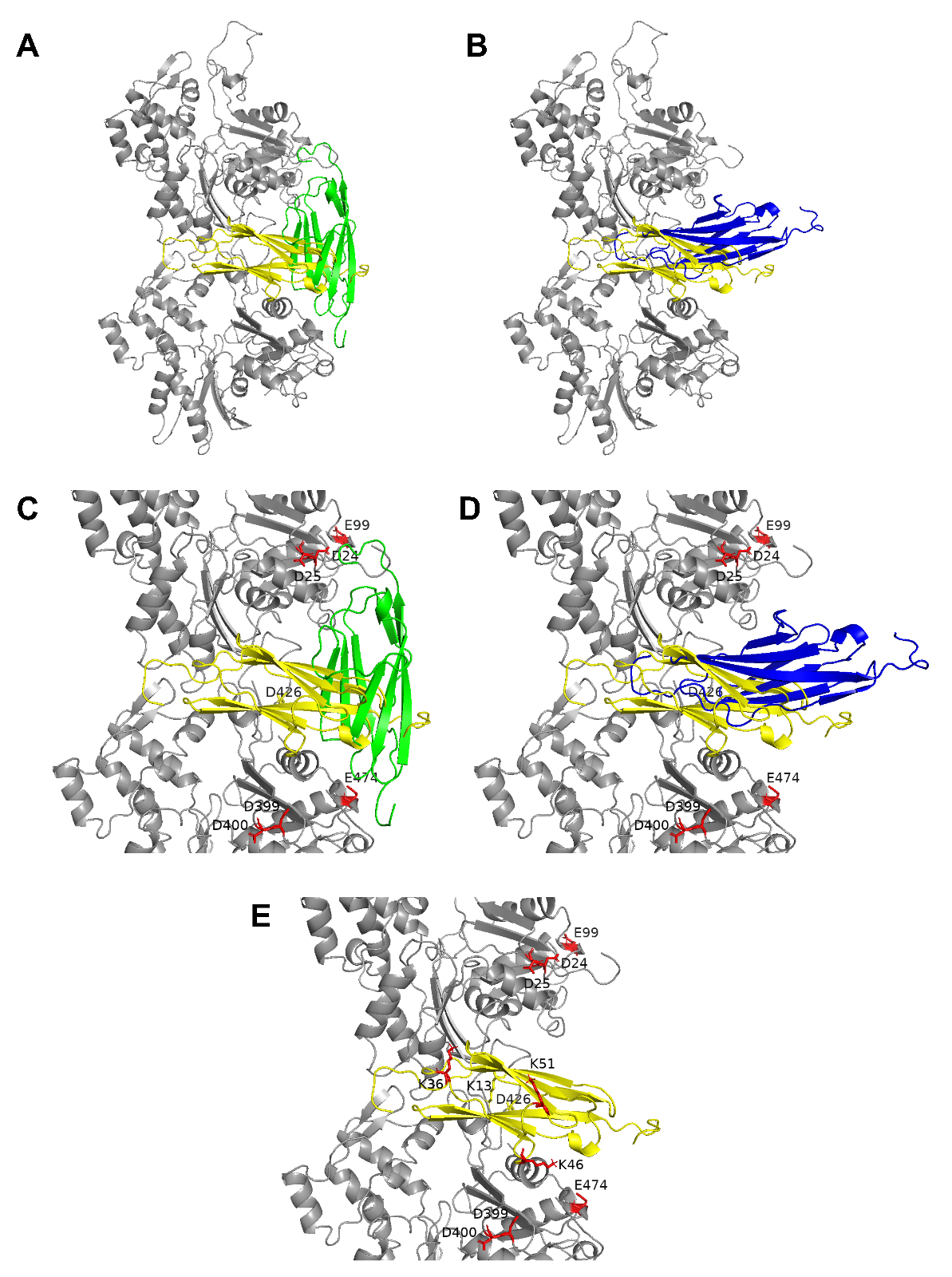

**Supplemental Figure S1: Top-scoring model from HADDOCK trial with no docking restraints and comparison to docking with XL-MS restraints.** (A, B) Docked Ig3 (yellow) to actin (grey) overlaid with structures docked with restraints that produced the highest initial HADDOCK (green) and Xwalk (blue) score, respectively. (C, D) Enlarged view with labeled actin residues and (E) enlarged view with labeled actin and Ig3 residues. For all structures, red residues were included in Pool 1 for HADDOCK trials and yellow residues were also included in Pool 2.

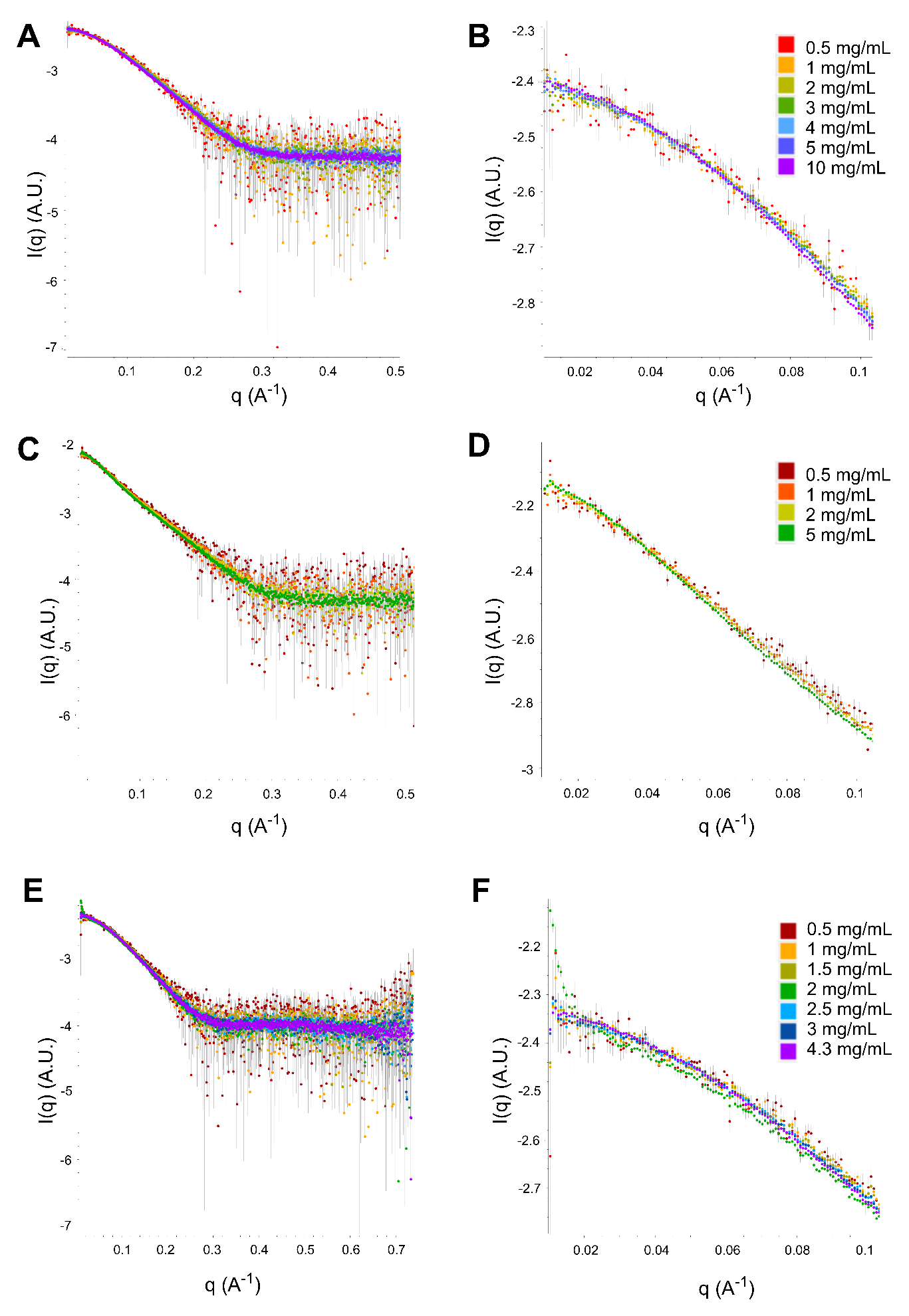

**Supplemental Figure S2: Overlay of buffer subtracted SAXS data for each palladin construct concentration series.** (A, B) Ig4, (C, D) Ig3-4, and (E, F) Ig3 scattering curves and low-q regions shown.

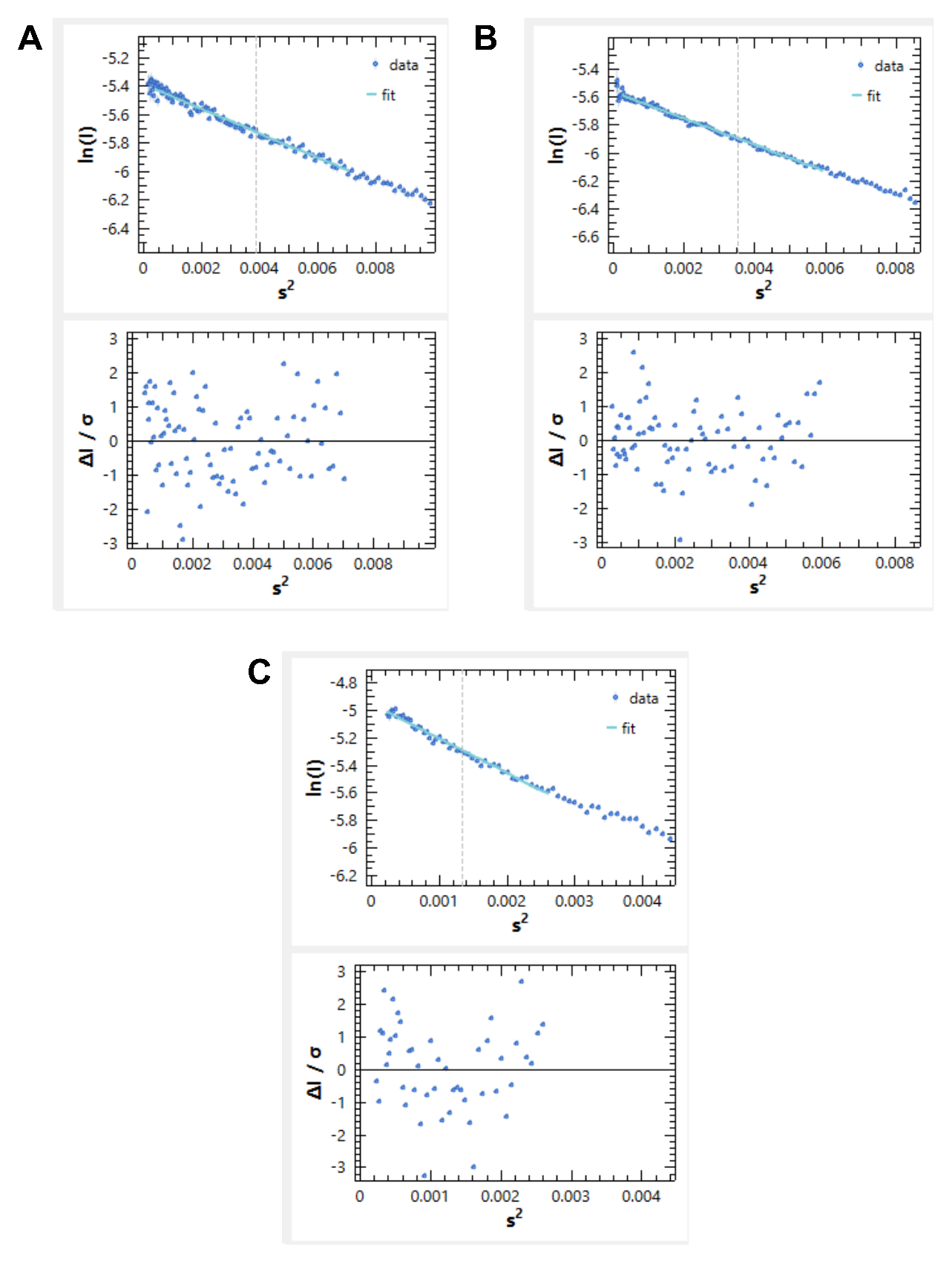

**Supplemental Figure S3: Guinier region and linear fit with residuals** for (A) Ig3 (B) Ig4 and (C) Ig3-4.

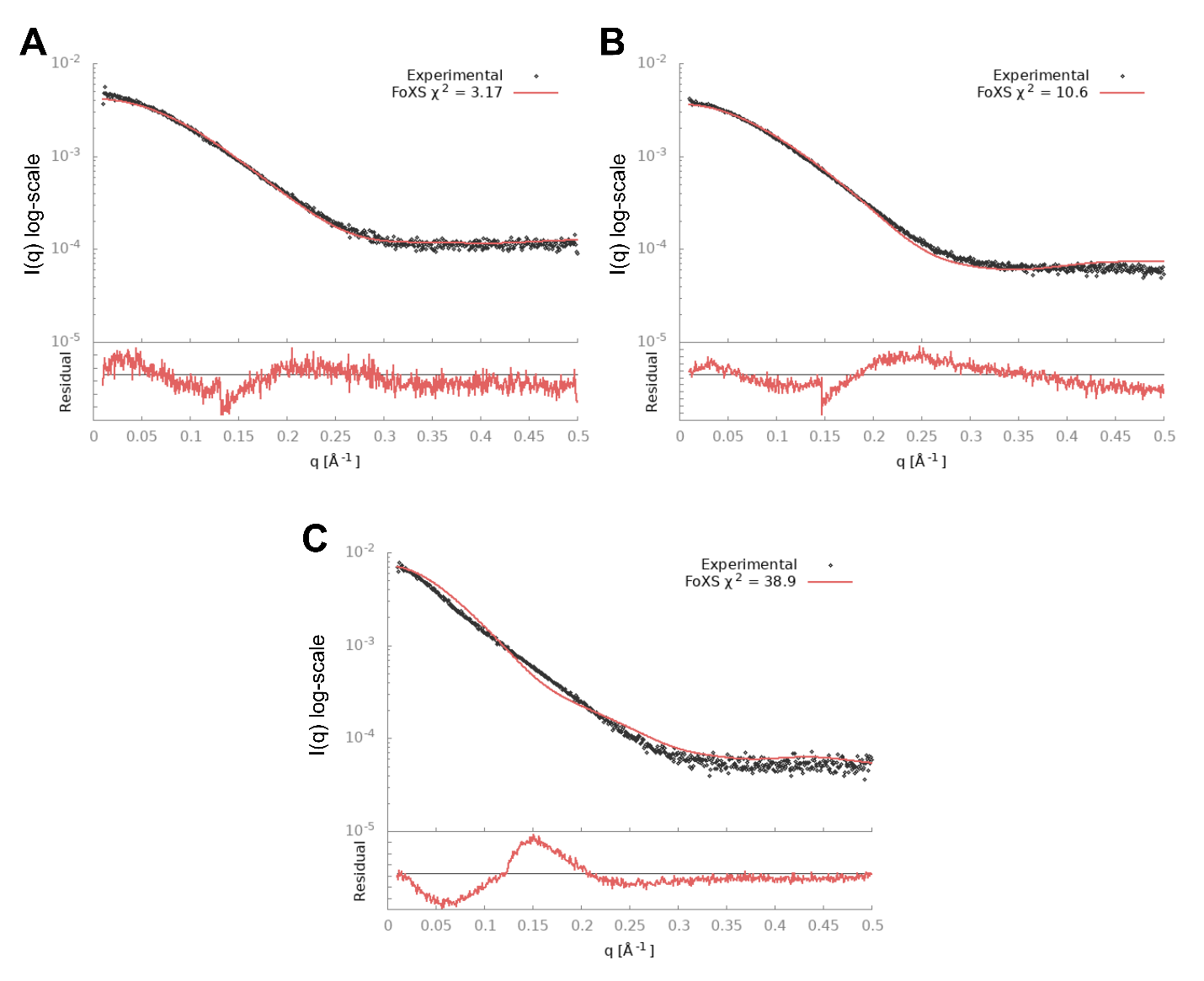
**Supplemental Figure S4: FoXS scattering curves with residuals and X2 values** for (A) Ig3 (B) Ig4 and (C) Ig3-4.

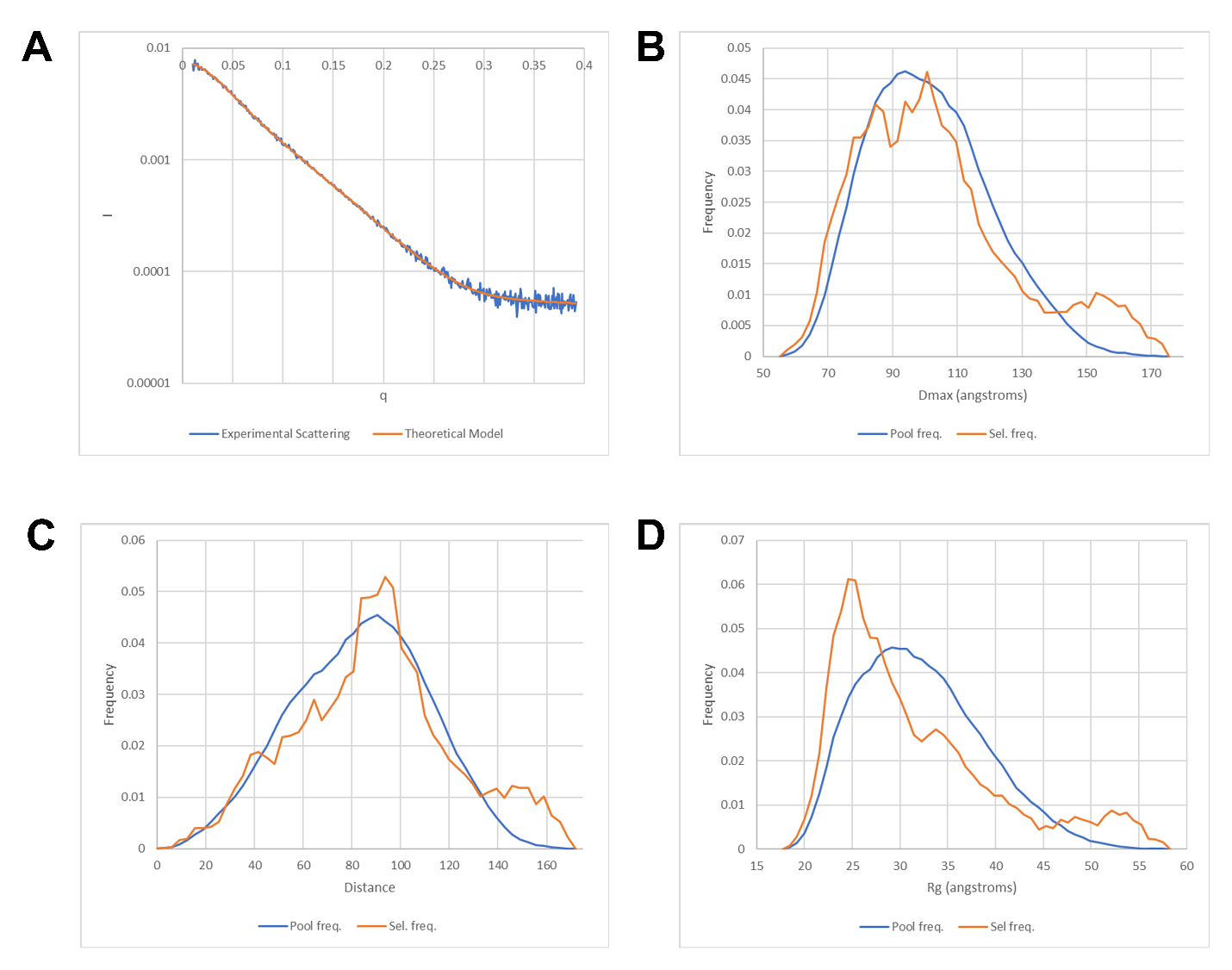

**Supplemental Figure S5: Fit and distribution plots of RANCH/GAJOE Ig3-4 modeling.** (A) goodness of fit between experimental SAXS scattering (blue) and theoretical scattering from RANCH/GAJOE model (orange). (B) D_max_ frequency of RANCH (blue) versus GAJOE selection (orange). (C) N-C distance frequency of RANCH (blue) versus GAJOE selection (orange) (D) Rg distance frequency of RANCH (blue) versus GAJOE selection (orange).
